# Transcriptomics based prediction of survival and response to therapy in malignant mesothelioma

**DOI:** 10.1101/2022.04.13.488207

**Authors:** Nishanth Ulhas Nair, Qun Jiang, Jun Stephen Wei, Vikram Alexander Misra, Betsy Morrow, Chimene Kesserwan, Leandro C. Hermida, Joo Sang Lee, Idrees Mian, Jingli Zhang, Alexandra Lebensohn, Manjistha Sengupta, Javed Khan, Eytan Ruppin, Raffit Hassan

## Abstract

Malignant mesothelioma is an aggressive cancer with limited treatment options and poor prognosis. Better understanding of mesothelioma genomics and transcriptomics could advance novel therapies. We performed whole-exome and RNA-sequencing of germline and tumors of 122 patients with pleural, peritoneal, and tunica-vaginalis mesothelioma. We identify a 48 gene prognostic signature that is highly predictive of mesothelioma patient survival including *CCNB1*, whose expression is highly predictive of patient survival on its own. Using a synthetic-lethality (SL) based pipeline for analyzing the patients’ transcriptomic data, we identified SL-based signatures predictive of response to an anti-PD1 immune checkpoint inhibitor and combination therapies with pemetrexed. These SL-profiles successfully predict the overall patient-response observed across targeted, immuno- and chemotherapies in 11 independent mesothelioma clinical trials spanning 7 different treatments. These findings lay a basis for future studies aimed specifically at testing the ability of these SL profiles to serve as treatment biomarkers in mesothelioma.

## INTRODUCTION

Malignant mesothelioma is an aggressive cancer arising from the mesothelial cell-linings of the pleura, peritoneum, pericardium, or tunica vaginalis with an annual incidence of 3,300 new cases in the United States (1). Malignant pleural mesothelioma (MPM) comprises 80% of the cases, while malignant peritoneal mesothelioma (MPeM) includes the remaining 15-20% (2). Pericardial and tunica vaginalis mesothelioma are very rare. Mesothelioma is a difficult to treat cancer with poor prognosis and limited treatment options. The median survival from diagnosis of MPM is approximately 12 months with treatment regimen of pemetrexed and cisplatin (3). Recently, FDA has approved the combination of immune checkpoint inhibitors, ipilimumab and nivolumab, as first-line therapy for MPM, with a modest but statistically significant increase in overall survival of 18.1 months over 14.1 months in the chemotherapy group (4). MPeM has a better prognosis than pleural mesothelioma, with median overall survival longer than 5 years, when amenable to cytoreductive surgery and hyperthermic intraperitoneal chemotherapy (2). Given the limited treatment options for patients who progress on pemetrexed based chemotherapy and checkpoint immunotherapy, there is an urgent need to develop new therapies.

Predisposing factors of mesothelioma are, exposure to asbestos fibers and prior radiation therapy (5-8). In addition, germline mutations in *BRCA1-*associated-protein 1 (*BAP1*) increase the risk of developing malignant mesothelioma as well as other common cancers (9). The somatic mutations that have been identified in MPM include loss-of-function mutations in tumor suppressor gene *BAP1*, as well as epigenetic regulatory genes *DDX3X* and *SETD2* (10). In addition, recurring deletions of chromosomal 3p21 (target gene *BAP1*), 9p21 (*CDKN2A*), and 22q12 (*NF2*) have been identified in malignant mesothelioma (11).

Since current therapies for mesothelioma lead to a good response only in a small set of patients (12-22), it is important to identify predictive biomarkers for patient response. Several prior studies with small number of patients have evaluated predictors of response to therapy and patient survival. These include MesoNet, a deep convolutional network approach, which uses whole-slide digitized images to predict the overall survival of mesothelioma patients (23). Using neural networks, a gene-expression based classifier was learnt from a small dataset of 21 MPM patients to predict survival (24). Another work used gene-expression ratio-based predictor derived from 17 mesothelioma tumors for determining treatment outcome (25). More recently, the loss of the tumor suppressor gene, *BAP1* has been proposed as candidate biomarker for immunotherapy treatment in mesothelioma (26).

A deeper knowledge of the genetic, transcriptomic, and immunogenic events involved in malignant mesothelioma is critical for successful development of prognostic biomarkers and personalized therapeutic modalities. In this study, we have three main aims. Our first goal is to present a new mesothelioma dataset of 122 patients with their genomic and transcriptomic profiles, as well as phenotypic and drug response information. Unlike previous large-scale studies that has focused on MPM patients (10,27), our dataset contains an approximately equal representation of MPM and MPeM patients, which allows us to identify differences between them. Our data is made publicly available, and it may be a useful resource for biologists and computational biology researchers interested in mesothelioma study. Our second goal is to come up with a transcriptome-based gene signature to predict mesothelioma patient survival in large-scale independent cohorts, and to help identify important genes which could possibly be potential drug target candidates. Finally, as a proof of concept, we apply a precision oncology framework which uses transcriptomic data and synthetic lethality (SL) predictions to predict drug response and suggest potential treatment options in mesothelioma patients. Our study may thus help to identify biomarkers for treatment-responsive subsets of malignant mesothelioma and provide guidance for novel clinical applications for mesothelioma therapy.

## METHODS

### Patients

All patients with malignant mesothelioma treated at the Thoracic Medical Oncology Clinic of the National Cancer Institute (NCI) were offered participation in a clinical trial of the natural history of malignant mesothelioma (ClinicalTrials.gov number NCT01950572). Between September 2013 and November 2019, 425 consecutive patients were enrolled, out of them 122 underwent whole exome sequencing of blood and tumor tissue. RNA-seq data was available for 100 patients. The NCI Laboratory of Pathology confirmed all diagnoses of mesothelioma and characterized their origin as pleural, peritoneal, or tunica vaginalis. Patients were enrolled regardless of asbestos exposure, age at diagnosis, or personal or family history of cancer. The study was conducted in accordance with the principles of the International Conference on Harmonisation - Good Clinical Practice (ICH-GCP) guidelines. Human subjects’ committees of the NCI approved the study and all patients provided written informed consent.

### Analysis of germline and somatic mutations and tumor mutational burden

Whole exome sequencing (WES) was carried out on paired tumor and germline DNA, which was extracted from formalin fixed paraffin embedded (FFPE) tumor tissue and from peripheral blood mononuclear cells (PBMC), respectively, as previously described (28,29). Exome sequencing was performed using Illumina NextSeq500 sequencers. The bcl files generated were converted to FASTQ files, which contained paired-end reads, using the bcl2fastq tool (Illumina, San Diego, CA). Germline variants found in the mesothelioma cohort (n = 122) were called using previously published methods (28,29). High-confidence germline variants were defined by the following criteria: total coverage of greater than 20x, Fisher score < 75 and variant allele frequency (VAF) ≥ 0.25. Germline variants were curated according to the American College of Medical Genetics and Genomics (ACMG) and the Association for Molecular Pathology (AMP) guidelines for the interpretation of sequence variants (30). Finally, pathogenic, or likely pathogenic tier 1 and 2 germline variants used in the Fisher’s exact test were filtered according to Phred score ≥ 20 and presence shown in Integrative Genome Viewer v. 2.3.31 (31). Somatic variant data were collected from VCF files of the genetic variants found in exome samples in tumors from the mesothelioma cohort (n = 122 tumors). The number of somatic mutations for each tumor were counted, and then divided by the number of bases in the exome of that tumor to yield the somatic mutation burden per MB.

### RNA-sequencing and data processing

RNA sequencing was performed on Illumina HiSeq2000 or NextSeq500 using TruSeq3 chemistry. The bcl files generated were converted to FASTQ files, which contained paired-end reads, using the bcl2fastq tool (Illumina, San Diego, CA).

*NCI mesothelioma data:* The RNA-seq reads were processed using the same workflow as the NCI Genomic Data Commons (GDC) has used to process TCGA-MESO RNA-seq reads, with the same settings where appropriate [see NCI GDC mRNA Analysis Pipeline in https://docs.gdc.cancer.gov/Data/Bioinformatics_Pipelines/Expression_mRNA_Pipeline/; (32)]. Namely, FASTQ reads were aligned to the NCI GDC reference genome GRCh38.p2 with GENCODE v22 gene annotations [see NCI GDC Reference Files in https://gdc.cancer.gov/about-data/gdc-data-processing/gdc-reference-files; (32)] using STAR v2.7.3a in two-pass mode. Read counts were quantified from STAR aligned read BAMs using HTSeq v0.11.2 in reverse strand-specific mode to reflect the RNA library protocol used. Counts were further processed for downstream analysis using edgeR with default settings (33). Genes with sufficiently large counts were filtered using the filterByExpr function, normalized using trimmed mean of M-values (TMM), and finally log counts-per-million (CPM) transformed all with default settings. There are 100 patients for which we had RNA-seq samples. TCGA mesothelioma data RNA-seq read counts were obtained from NCI GDC (27). Similar processing as above was done for the TCGA dataset. TCGA pan-cancer RNA-seq TPM (log transformed) data was taken from the Xena browser (34).

### Protein complex-enrichment and cluster gene expression

Mesothelioma prognostic signature gene-set was used for protein complex-enrichment analysis. Protein complex dataset used is 03.09.2018 Corum 3.0 current release. Fisher exact test was used for the enrichment studies (FDR < 0.05). Hierarchical clustering of the 48 genes composing the gene mesothelioma prognostic signature was done using the hclust function in *R* (https://stat.ethz.ch/R-manual/R-devel/library/stats/html/hclust.html).

### Immune cell abundance estimates

CIBERSORT was used to estimate the immune cell abundance in each patient in the NCI mesothelioma dataset, and also in the TCGA and Bueno *et al*. mesothelioma patient data sets. CIBERSORT was run for 500 permutations using the default gene signature LM22 CIBERSORT. The results were run in the ‘Relative’ mode generating ‘Relative’ fractions of 22 immune cell types for each of the 100 patients in the NCI mesothelioma dataset. The relative values indicate the relative abundance of each immune cell type as a fraction of the bulk tumor data (which does not include cancer). The relative values for each sample add up to 1 for all cell types. The immune estimates for TCGA cancer patients were also obtained using a CIBERSORT analysis from (35). For all the immune cell types which have more than 5% mean relative abundance across samples in the NCI mesothelioma dataset, we tested their association with patient survival (FDR < 0.2).

### Coverage analysis using synthetic lethality and synthetic rescue interactions

SELECT (SynthEtic Lethality and rescue-mediated precision onCology via the Transcriptome) is a computational method to stratify patients as responders or non-responders for each drug using whole transcriptomic data (36). SELECT first infers clinically-relevant synthetic lethal (SL) or synthetic rescue (SR) pairs across cancer types by mining thousands of patient tumor genomic and transcriptomic data in the TCGA dataset. For each of the different cancer therapies, we mapped the drugs to the targets (which they inhibit) using DrugBank (37). For chemotherapy and targeted therapy drugs, for each patient sample, SELECT assigns a risk score – called SL-score – based on the number of downregulated SL partners of the target genes inhibited by a drug. The method works under the assumption that a drug is likely to kill a tumor more effectively when its SL partners are downregulated, as the inhibition of the target by the drug will lead to active SL formations (as both the SL gene pairs will becoming jointly downregulated). For immunotherapy drugs, SELECT assigns risk scores using SRs and it works under the assumption there is an innate resistance to a drug if there are a lot of SR partners that are downregulated in the cancer sample, as the inhibition of the target by the drug will lead to active SR formations. For both SL and SR based approaches, higher the risk score, more likely the patient would respond to a drug (36). These risk scores are then used for drug response validations and classifying patients as responders or non-responders in many independent datasets. Following the findings of (36), a risk score > 0.44 was used to classify a patient as responder for chemotherapy or targeted therapy and a risk score ≥ 0.9 was used to classify a patient as responder for immunotherapy. We emphasize that our analysis uses the exact same parameters and thresholds that was used in the original SELECT pipeline (36), to identify the pertaining SL and SR partners and to compute risk scores in the mesothelioma cohorts, without any training and parameter tuning on the latter.

### Statistical analysis

For survival analysis, survival data was censored as of November 1, 2019. The K-M method was used to estimate the survival probability from diagnosis date to death or last follow-up (censored) date. Data were stratified either by site (peritoneal, pleural) alone, or were first analyzed according to subgroups of asbestos, smoking history, diagnosis age, and sex, and then stratified by site. Hazard ratios (and their 95% CI) were calculated by Cox proportional hazard model with pleural mesothelioma as reference. Median overall survival estimates were provided with their 95% CI (however, some were non-estimable). K-M plots are presented with pointwise 95% confidence limits and the number at risk for various points. As some of the mesothelioma samples in the NCI mesothelioma data were acquired post treatment, we computed the survival of all patients from the time of biopsy sampling to the time of death or last follow-up. This survival time estimate was used for the gene expression and copy-number based survival association studies.

Mesothelioma prognostic signature set identification and cross-validation was done based on survival analysis. Risk scores for each patient is computed as the median expression of the mesothelioma prognostic signature gene-set for that patient. For the NCI mesothelioma data, we did the analysis using a “leave-one-out” cross validation. For each of the training samples, we re-computed the genes whose increased expression is associated with worse survival using Cox regression analysis after controlling for age, sex, and site-of-disease (FDR < 0.1), and the median expression value (risk score) for this set of genes was computed for the test data point. The risk scores were then associated with survival analysis using Cox regression (after controlling for age, sex, and site of disease).

To calculate differential gene expression between pleural and peritoneal mesothelioma, we used edgeR (33) on the RNA-seq count data to compute differential expression between the two groups, after controlling for various confounding factors like age and sex (FDR < 0.1).

## RESULTS

### NCI mesothelioma dataset and patient characteristics

We performed whole exome sequencing (WES) of paired blood and tumor tissues from 122 malignant mesothelioma patients who participated in the clinical trial at NCI (ClinicalTrials.gov number NCT01950572) to detect somatic and germline mutations. Patient population was enriched for those who carried a previously described germline mutation in BROCA V10 panel target genes (38). Our study cohort includes an equal proportion of patients with pleural and peritoneal mesothelioma, as compared to published mesothelioma cohorts which only included patients with pleural mesothelioma (10,27). We also performed RNA-seq analysis on 100 tumor samples for which RNAs were available (Methods). Phenotypic characteristics and drug response information is also presented. This data resource is made publicly freely available in https://clinomics.ccr.cancer.gov/clinomics/public/ and in **Table 1** and **Tables S1-S4**.

**Table 1:** Mesothelioma patient characteristics.

| **Patient characteristics** | ***n* (%)** |
| --- | --- |
|  | 122 (100) |
| Sex |  |
| Male | 69 (57) |
| Female | 53 (43) |
| Age at diagnosis, y |  |
| <60 | 76 (62) |
| ≥60 | 46 (38) |
| Asbestos exposure* |  |
| Yes | 67 (54) |
| No | 33 (27) |
| Unknown | 22 (19) |
| Mesothelioma site |  |
| Pleura | 59 (48) |
| Peritoneum | 61 (50) |
| Tunica Vaginalis | 2 (2) |
| Radiation prior to mesothelioma Dx |  |
| Yes | 9 (8) |
| No | 113 (92) |
| Prior Mesothelioma treatment |  |
| None | 69 (57) |
| Platinum based chemo | 40 (33) |
| Chemo plus immunotherapy | 10 (8) |
| Immunotherapy | 2 (2) |

\*Asbestos exposure by self-report

Clinical features and characteristics of patients in the NCI mesothelioma cohort are shown in **Table 1**. Majority of patients are male (57%) with average age at diagnosis being 54.2 years (range 12 to 80 years). Large proportion of patients were diagnosed early (62% <60 years old). Peritoneal (50%) and pleural (48%) mesothelioma cases existed in almost equal proportions. Most tumors had epithelial histology (84%); average age at diagnosis of patients with pleural mesothelioma was higher than that of peritoneal mesothelioma (58.9 + 10.9 years vs 50.0 + 14.4 years, P<0.0001) and were more likely to self-report asbestos exposure (61% vs 47.5%, P = 0.02). Patients in our study were largely treatment naïve at the time of tissue sequencing (57%), or had received platinum-based chemotherapy (33%), with a small proportion of patients who received immunotherapy (2%), or a combination of chemo and immune therapy (8%).

### Germline and somatic mutations in mesothelioma

In the NCI mesothelioma cohort of 122 patients (n = 59 for MPM, n = 61 for MPeM, n = 2 for tunica vaginalis mesothelioma), whole exome sequencing (WES) analysis of germline variants was performed according to the American College of Medical Genetics and Genomics (ACMG) and the Association for Molecular Pathology (AMP) guidelines for variant interpretation (30) (**Fig. 1**). A total of 43 Pathogenic (P) or Likely Pathogenic (LP) variants in 21 cancer predisposition genes were detected in 37 patients **(Table S1)**. Consistent with previous studies, *BAP1* P/LP variants were the most common alterations seen in this cohort (13.1%), with similar rate of *BAP1* P/LP in MPM (11.9%) and MPeM patients (14.8%). P/LP variants in *BAP1* were found in 16 patients, accounting for 37% of the total P/LP variants seen in this cohort. Of these 16 patients with heterozygous P/LP *BAP1* variants, 3 patients were also found to be heterozygous carriers for a second P/LP variant in a gene predisposing to an autosomal recessive (AR) cancer syndrome (*ERCC2, SBDS, and XPA*) and 1 patient was heterozygous for a pathogenic variant in *SDHA*, associated with an autosomal dominant predisposition to Paragangliomas. Eight additional patients were each positive for a P/LP variant in an AD cancer predisposition gene (*ATM, BRCA1, BRCA2, DDX41, MLH1, POT1, TP53, WT1*).

**Figure 1:**
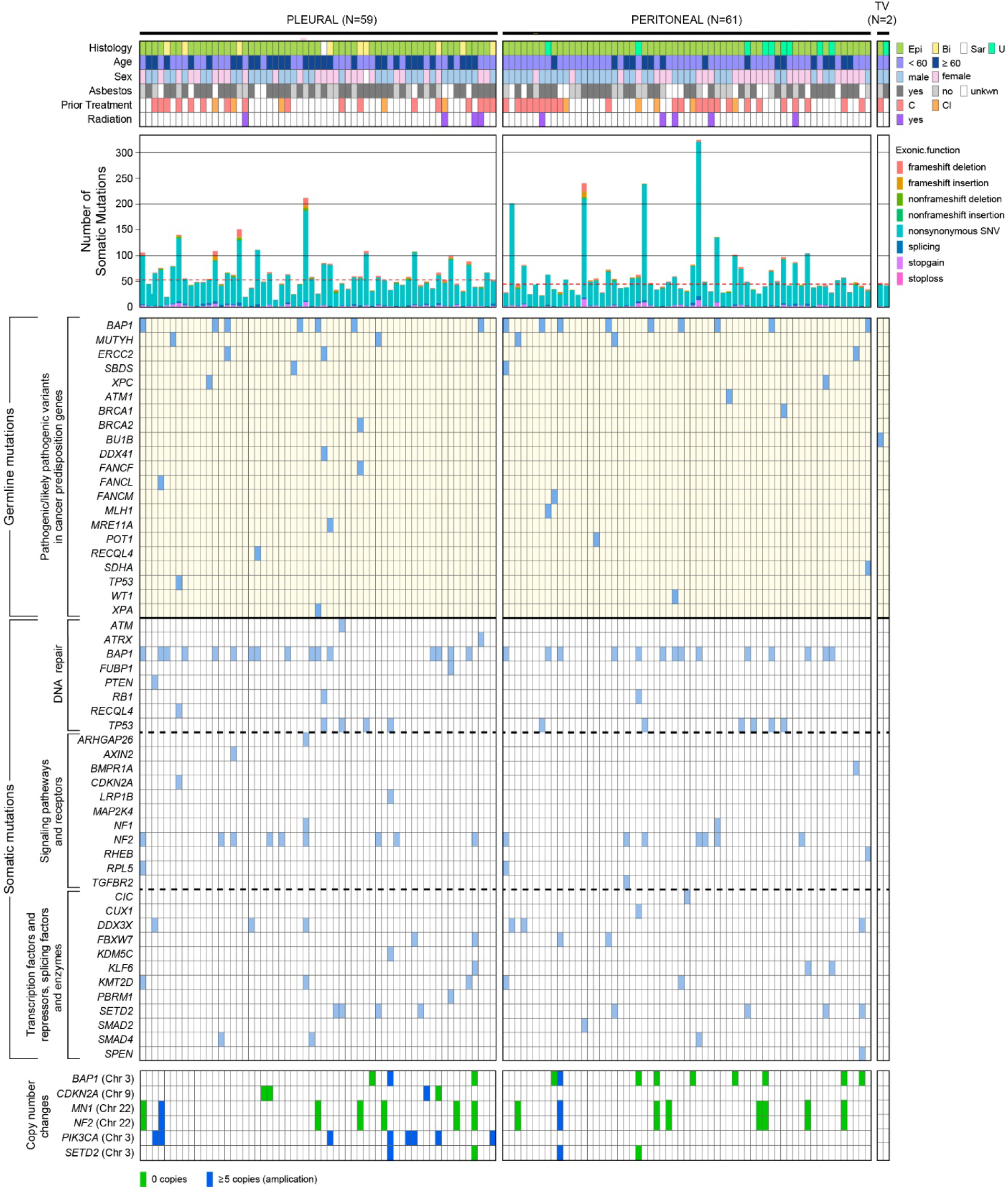
Overview of the NCI mesothelioma data and mutational signature analysis. Demographic information, tumor histology and mutational profiles of patients with mesothelioma. The mesothelioma cohort in this study is comprised of 122 patients, including patients with pleural (n = 59), peritoneal (n = 61) and tunica vaginalis (n = 2) forms of mesothelioma. The mutational profile consists of germline mutations in pathogenic and likely pathogenic cancer predisposition genes, as well as somatic mutations in genes associated with signaling pathways and receptors, DNA repair genes, and transcription factors and repressors, splicing factors, and enzymes. Copy number changes are indicated in relevant genes.

For somatic mutations (also shown in **Fig. 1)**, the median number of mutation burden in a tumor is 53 mutations in MPM (a burden of 1.17 mutations/MB, similar to what has been reported in mesothelioma (39,40), 45 mutations in MPeM (1.00 mutations/MB), and 44.5 mutations (0.98 mutations/MB) in tunica vaginalis mesothelioma. Indels were more common in the pleural form than in the peritoneal form. The genes that most frequently carried somatic mutations include *BAP1* (24.59%), *NF2* (13.11%), *TP53* (8.2%), *SETD2* (6.56%). No significant difference in the frequency of these gene mutations were observed between MPM and MPeM. We also performed K-M (41) analysis to compare the overall survival times between MPM and MPeM (**Fig. S1**). In general, MPeM patients showed a higher overall survival than MPM patients in our cohort (log-rank test P = 0.043). We then analyzed the copy-number variations (**Table S2**). *BAP1* copy number loss was observed less in MPM (3.39%) than MPeM (13.11%), while *PIK3CA* amplification was observed only in MPM (13.56%) but not MPeM.

### Identifying a gene expression signature associated with survival in mesothelioma

We next analyzed gene expression based on RNA-seq data for 100 NCI mesothelioma patients (**Table S3**) and identified the genes associated with patient survival. Since some of the mesothelioma samples in the NCI mesothelioma data were acquired post treatment, we recomputed the survival of all patients from the time of biopsy sampling to the time of death or last follow-up (**Table S4a**) and used this survival data for our gene expression and copy-number based survival association studies.

We identified 48 genes in the NCI mesothelioma data, whose increased expression is associated with worse survival using Cox regression analysis, after controlling for age and sex (FDR < 0.1). We term this set of 48 genes the “*mesothelioma prognostic signature*” gene-set and it is used for further analysis (**Table S5a, Supp. results**). We also identified 27 genes with increased expression that were associated with better survival (Cox regression analysis, after controlling for age and sex, FDR < 0.1; **Table S5b**). We did not find any genes whose copy number variation was associated with mesothelioma patient survival (after controlling for age and sex, FDR < 0.1). Gene ontology (GO) enrichment analysis on the mesothelioma prognostic signature gene-set showed a strong enrichment for GO terms related to cell cycle processes, DNA repair, chromosome organization, telomere organization, proliferation, gene silencing, and others. (**Fig. 2a, Table S5c**).

**Figure 2:**
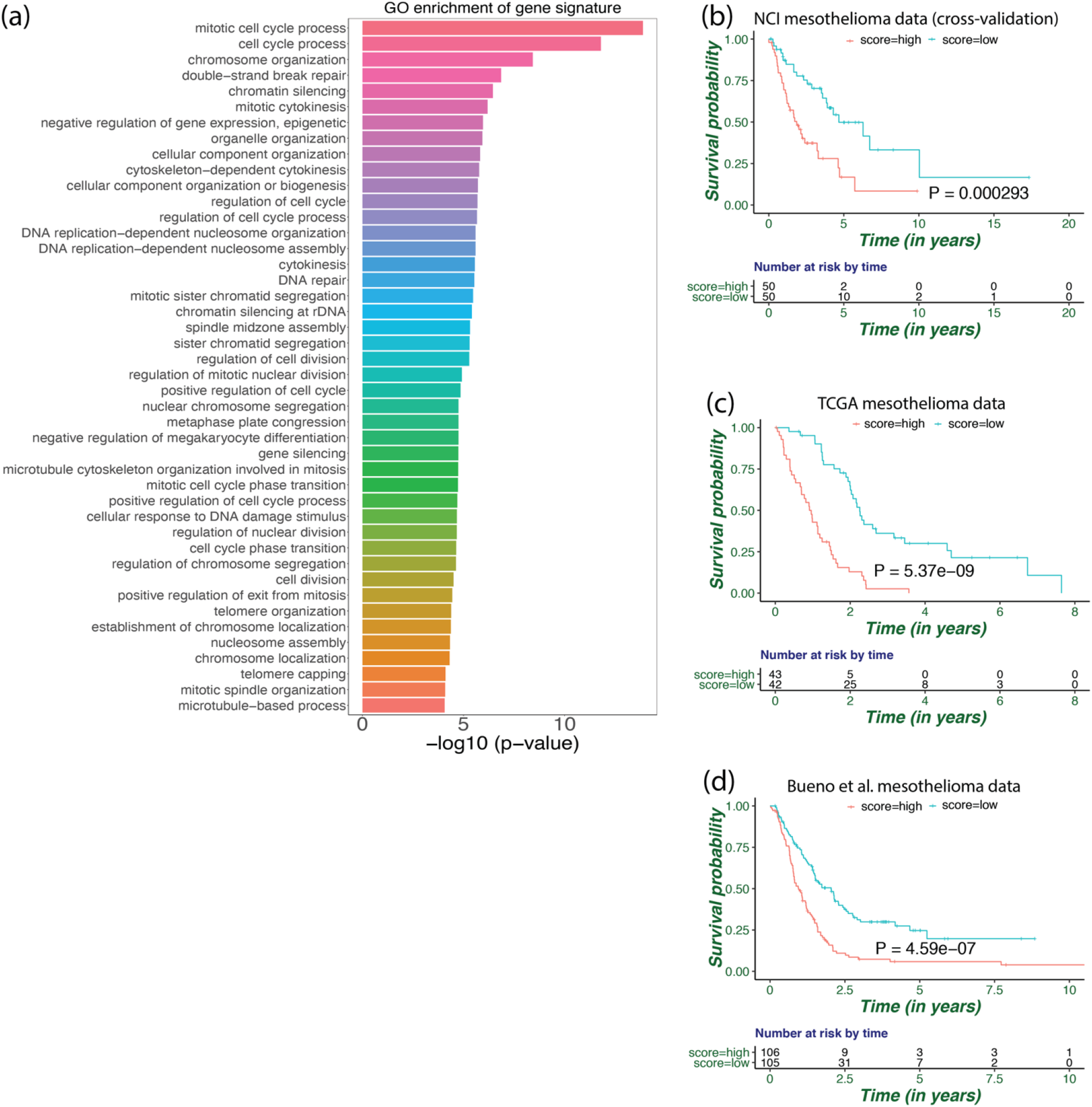
Gene ontology enrichment analysis and validation of mesothelioma prognostic signature. **(a)** Gene ontology (GO) enrichment analysis of the mesothelioma prognostic signature using GOrilla tool. GO terms with p-values < 1e-4 are shown here (list of all GO terms with p-values < 1e-3 are enlisted in **Table S5c**). **(b-d)** Survival analysis using K-M plot of patients in top 50 (high risk) and bottom 50 (low risk) percentile based on their predicted risk scores, in **(b)** NCI mesothelioma dataset in cross validation, **(c)** TCGA mesothelioma dataset, and **(d)** Bueno et al. mesothelioma dataset. Log-rank test p-values are shown.

### Mesothelioma prognostic signature is highly predictive of mesothelioma patient survival in other independent patient cohorts

We aimed to validate if the expression of the mesothelioma prognostic signature genes could predict patient survival in independent mesothelioma patient cohorts (RNA-seq data was processed as TMM log CPM format, Methods). We defined a *combined risk score* as the median expression of the 48 genes in the mesothelioma prognostic signature set and computed it for each patient. Since the increased expression of these genes were associated with worse survival, we hypothesize that their risk scores would be a marker for survival in mesothelioma patients. Using Cox regression analysis in cross-validation (Methods), after controlling for age, sex, and site-of-disease, we found, as expected, that high risk scores were indeed associated with worse survival outcome in the NCI mesothelioma data (Hazard Ratio or HR = 1.76, P = 6.87e-04, **Fig. S2, Table S5d**). A K-M survival analysis also showed similar results (log-rank test P = 0.00029, **Fig. 2b**).

To test the predictive value of the mesothelioma prognostic signature, we validated it in two large, independent, mesothelioma datasets: the TCGA mesothelioma dataset of 85 pleural mesothelioma patients (27) and Bueno *et al*. dataset of 211 pleural mesothelioma patients (10). We computed the risk scores using the mesothelioma prognostic signature for each patient in these datasets and predicted overall survival. Cox regression analysis controlling for age and sex showed that increased risk scores was associated with worse patient survival in both TCGA mesothelioma (HR = 2.6, P = 6.94e-10, **Fig. S2**) and Bueno *et al*. dataset (HR = 1.49, P = 4.34e-07, **Fig. S2**). Similar results were obtained using a K-M analysis (log-rank test P = 5.37e-09 for TCGA, **Fig. 2c**; and log-rank test P = 4.59e-07 for Bueno *et al*. data, **Fig. 2d**). Control experiments were performed using randomized gene sets of same size as those present in mesothelioma prognostic signature gene-set, and risk scores were computed. Reassuringly, these random control experiments did not show any survival association (1000 iterations, randomization test empirical P < 0.001 while comparing the actual HRs with the control; see **Fig. S3** for details). The list of 27 genes whose increased expression is associated with better survival did not show a survival association in the TCGA dataset and showed only some significant survival association in the Bueno *et al*. data set (**Fig. S4**).

To test the robustness of predicting survival using the 48-gene mesothelioma prognostic signature, we computed a risk score using an alternative method. For each patient, a *fractional risk score* was computed by counting the fraction of the 48 genes in the mesothelioma signature which were highly expressed (top 33 percentile of all genes) in a patient. We found this score to be a strongly associated with patient survival confirming our findings (results are quite similar to using computing combined risk scores; explained in **Supp. results, Fig. S5**).

We checked the association of gene expression with survival for TCGA mesothelioma and Bueno *et al*. datasets separately, using Cox regression after controlling for age and sex. For the TCGA mesothelioma and Bueno *et al*. datasets, there are 2144 and 1210 genes respectively, whose increased expression is significantly associated with decreased survival (FDR < 0.1). We found that the mesothelioma prognostic signature of 48 genes have significant overlap with the survival associated genes from the TCGA (P = 9.41e-15) and Bueno *et al*. mesothelioma (P = 1.85e-14) datasets (using a hypergeometric test, see **Supp. results** for details). There are 31 over lapping genes between the three studies whose increased expression is significantly associated with decreased survival in all three mesothelioma datasets (FDR < 0.1, **Table S5e, Supp. results**). Some of these 31 genes could be potential candidates for drug targeting (provided they are druggable), as inhibiting them could potentially improve patient survival (**Table S5e**). GO enrichment analysis on these 31 genes shows a strong enrichment for GO terms related to various metabolic processes (**Table S5f**).

### *CCNB1* gene is strongly predictive of mesothelioma patient survival

We explored the possibility of identifying a smaller subset of genes among the mesothelioma prognostic signature genes that could possibly play an important functional role in mesothelioma, with the hope that some of these genes may be potential candidates for targeted therapies. Previous work by Melaiu *et al*. has identified 51 genes that were differentially upregulated consistently across various studies in mesothelioma versus non-malignant mesothelial samples (42). Two of these genes (*CCNB1* and *NUSAP1*) were present in our 48-gene mesothelioma signature gene set.

We calculated the risk factors associated with the increased median expression of these 51 genes and found them to be predictive of worse mesothelioma patient survival, both in the TCGA and Bueno *et al*. mesothelioma cohorts (Cox regression after controlling for age and sex; HR = 1.70, P = 9.91e-05 for TCGA and HR = 1.37, P = 4.96e-04 for Bueno *et al*. mesothelioma data; **Fig. S6**). The median expression of *CCNB1* and *NUSAP1* genes, by themselves is predictive of patient survival as well (Cox regression after controlling for age and sex; HR = 2.28, P = 1.67e-07 for TCGA and HR = 1.40, P = 2.37e-05 for Bueno *et al*. mesothelioma data; **Fig. S6**). Among these two genes, we found that the increased gene expression of *CCNB1* alone was highly associated with worse mesothelioma patient survival outcome (Cox regression after controlling for age and sex; HR = 2.54, P = 1.89e-08 for TCGA mesothelioma and HR = 1.40, P = 1.65e-05 for Bueno *et al*. mesothelioma data; **Fig. S6**). Although these survival associations are not as strong as that those obtained using the 48-gene mesothelioma prognostic signature, they are still notable.

### The protein-protein interaction (PPI) network connecting mesothelioma prognostic signature genes support the functional importance of CCNB1 and the complexes it forms

To further explore the pathways in which the 48-genes in “mesothelioma prognostic signature” are involved, we looked at the protein-protein interactions among them. The PPI network for these genes were inferred from the STRING dataset and website, for the 48 genes (43). We identified three different cliques in the network – clique 1 with 21 genes, clique 2 with 5, and clique 3 with 6 genes (**Fig. 3a**). We suggest that high degree hub proteins (i.e., proteins with a lot of interacting partners in the PPI network) in these cliques could play a functional role in mesothelioma and could be potential drug targets. We note that both *CCNB1* and *NUSAP1* are such hubs in clique 1. *CCNB1* and *NUSAP1* have degree 19 (ranked 2 out of 48) and 16 (ranked 6 out of 48) respectively in the PPI network (degree of a protein is the number of interacting partners for that protein in the PPI network). *MCM2* with degree 22 has the highest rank.

Protein complex enrichment analysis was performed on clique 1 genes, which forms the largest clique in the PPI network discussed above, using the CORUM protein complex resource [Methods, (44)]. These genes are enriched for 2 complexes – CCNB1-CCNF and CDK1-CCNB1-CCNF complex (Fisher test, FDR < 0.05, **Fig. 3b, Table S5g**), in which *CCNB1* and *CCNF* are members. These complexes functional annotation indicates that they are involved in cell cycle and DNA processing.

**Figure 3:**
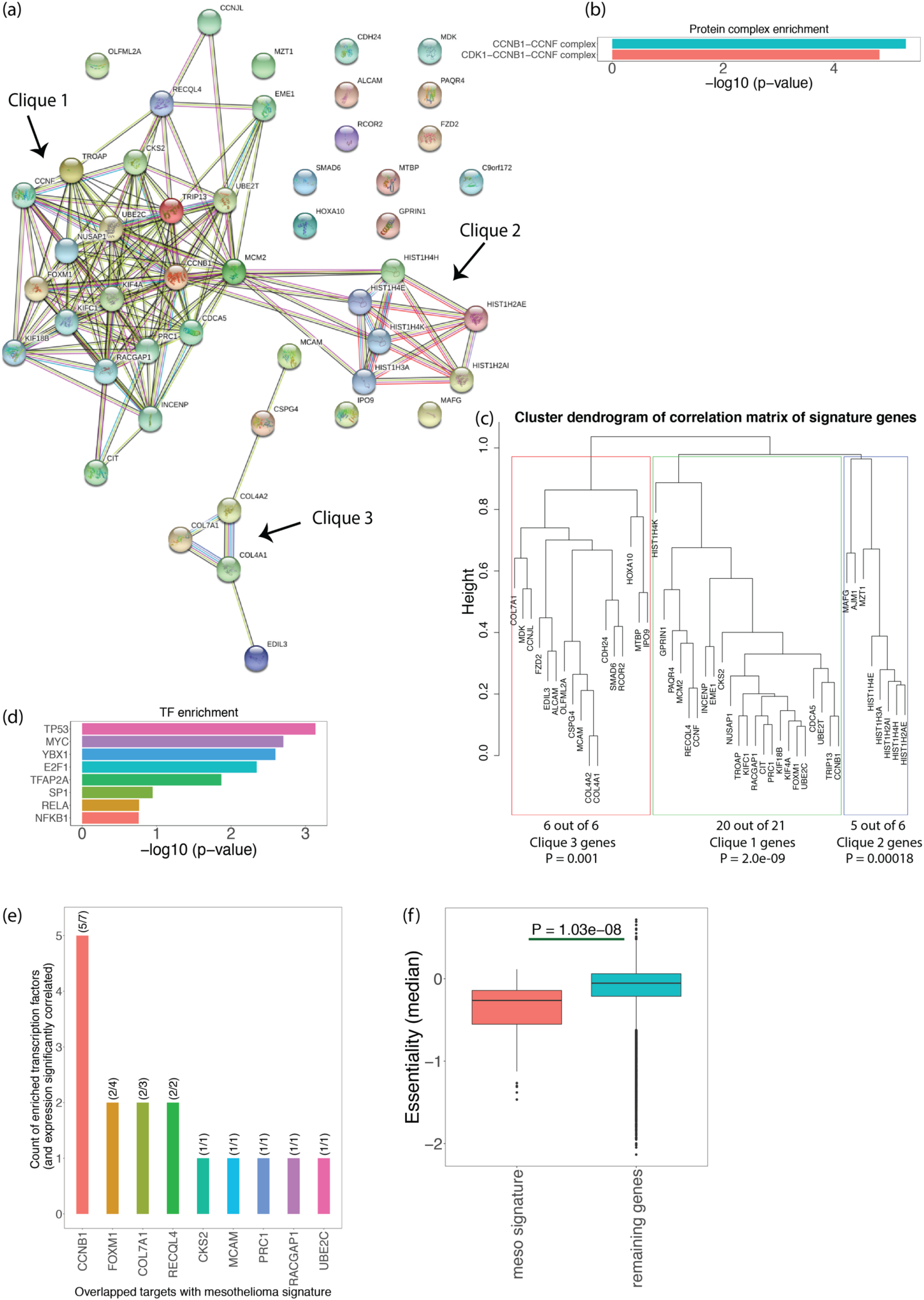
Mesothelioma prognostic signature analysis. **(a)** Protein-protein interaction (PPI) network of the genes in the mesothelioma prognostic signature identified 3 cliques in the NCI mesothelioma dataset by STRING. **(b)** Protein complex enrichment analysis of Clique 1 genes. These genes are enriched in 2 complexes, CCNB1-CCNF complex and CDK1-CCNB1-CCNF complex. Fisher exact test, FDR < 0.02). X-axis represents Fisher test negative log_10_ p-values. **(c)** Correlation matrix was computed on the matrix of 48-gene mesothelioma prognostic signature across all patients. Hierarchical clustering Identified three clusters that significantly overlapped with the three cliques obtained in the PPI network. Fisher exact test p-values are shown. **(d)** Mesothelioma prognostic signature is enriched for 8 transcription factors (TFs) (details provided in **Table S5h**). **(e)** Corelation between expression of mesothelioma signature genes and corresponding TFs. Expression of every gene in the mesothelioma prognostic signature cohort that is potentially regulated by one or more of the 8 enriched TFs shown in (d), was correlated with the expression of the corresponding mapped TF (FDR < 0.2). The number of such enriched TFs which were significantly correlated (FDR < 0.2) with these genes is plotted as a bar graph. The count of enriched TFs that are significantly correlated vs the total number of TFs mapped to a particular gene is shown in parenthesis. **(f)** Median essentiality values between from CRISPR-Cas9 gene-knockout essentiality screens (from 7 pleural mesothelioma cell lines (ACCMESO1, NCIH2452, NCIH2052, MPP89, ISTMES2, MSTO211H, NCIH28) for each gene is computed, and compared between genes in the mesothelioma prognostic signature vs the remaining genes. (Less essentiality value implies the gene is more essential.) One-sided Wilcoxon rank-sum test p-value is shown.

We computed the correlation between the expression of the 48-genes present in mesothelioma prognostic signature, across all patients and performed a cluster analysis on their expression patterns (Methods). Again, three clusters were identified that significantly overlapped with the respective cliques obtained in the PPI network (**Fig. 3c**, Fisher exact test p-values shown), together providing convergent evidence to the functional importance of these genes and these two complexes.

### A transcription factors enrichment analysis further points to the central role of *CCNB1*

We then aimed to identify the transcription factors that may control the expression of the mesothelioma prognostic signature genes, using the TRRUST v2 package (45) for transcription factor (TF) enrichment analysis. The results are summarized in **Table S5h**. We found significant enrichment for 8 TFs – *TP53, MYC, YBX1, E2F1, TFAP2A, SP1, RELA and NFKB1* (**Fig. 3d, Table S5h**), which together regulate 9 genes of the 48 composing the mesothelioma prognostic signature (**Fig. 3e, Table S5h**). Notably, out of these 8 TFs, 7 have biding sites and regulate *CCNB1*, again pointing to its putative central functional role in mesothelioma. **Fig. 3e** portrays the number of signature genes whose expression is correlated with the corresponding TF gene expression (Spearman correlation, FDR < 0.2). This may provide additional evidence that these TFs potentially regulate these genes. Notably, *CCNB1* expression is significantly correlated with 5 out of these 7 TFs.

### Mesothelioma prognostic signature genes tend to be essential in CRISPR screens of mesothelioma cell lines

Since we are interested in investigating a subset of the 48-gene mesothelioma prognostic signature which could be potential drug candidates, we analyzed CRISPR-Cas9 gene-knockout essentiality screens (46,47) from 7 pleural mesothelioma cell lines (ACCMESO1, NCIH2452, NCIH2052, MPP89, ISTMES2, MSTO211H, NCIH28). We took the median essentiality values across the 7 cell lines for each gene. We found that the mesothelioma prognostic signature genes tend to be more essential than the rest of the genes (one-sided Wilcoxon rank-sum test P = 1.03e-08, difference in median essentiality score = 0.21, **Fig. 3f**). **Table S5i** shows the essentiality values for the signature genes. Ten genes (*MTBP, PRC1, INCENP, MCM2, RACGAP1, IPO9, CDCA5, KIF4A, CCNB1, KIF18B*) have essentiality in the top 10 percentile of all genes in these cell lines. *MTBP* is the top essential gene among the 48-gene signature (ranked 98.7 percentile among all genes) and *CCNB1* is also essential (ranked 91.6 percentile among all genes).

### Immune cell type abundance estimates show differential abundance of certain immune cell types in patients with mesothelioma

Next, we looked for immune cell types that were relatively high in mesothelioma, with the aim to understand mesothelioma immune microenvironment. We used CIBERSORT (48) to estimate the abundance of 22 immune cell types in each patient in the NCI mesothelioma dataset (Methods, **Table S6**). The estimates of immune cells of TCGA cancer patients were also obtained using a CIBERSORT analysis from Lee et al, 2019 (35). The relative immune cell abundance estimates were compared between the NCI mesothelioma dataset (NCI MESO), TCGA mesothelioma patients (TCGA MESO), TCGA pan-cancer patients (mesothelioma not included), TCGA lung adenocarcinoma (TCGA LUAD) and lung squamous cell carcinoma (TCGA LUSC) (**Fig. 4a)**. Some immune cell types like, CD4+ memory resting T-cells, naïve B cells, activated Mast cells and monocytes, had higher relative abundance in NCI mesothelioma dataset, than in other cancer types in the TCGA (Wilcoxon rank-sum test, FDR < 0.1, **Fig. 4a**). We also observed that some immune cell types, like CD8+ T cells, macrophages M0, have lower abundance in NCI mesothelioma dataset than in other cancer types in the TCGA data base (Wilcoxon rank-sum test, FDR < 0.1, **Fig. 4a**).

**Figure 4:**
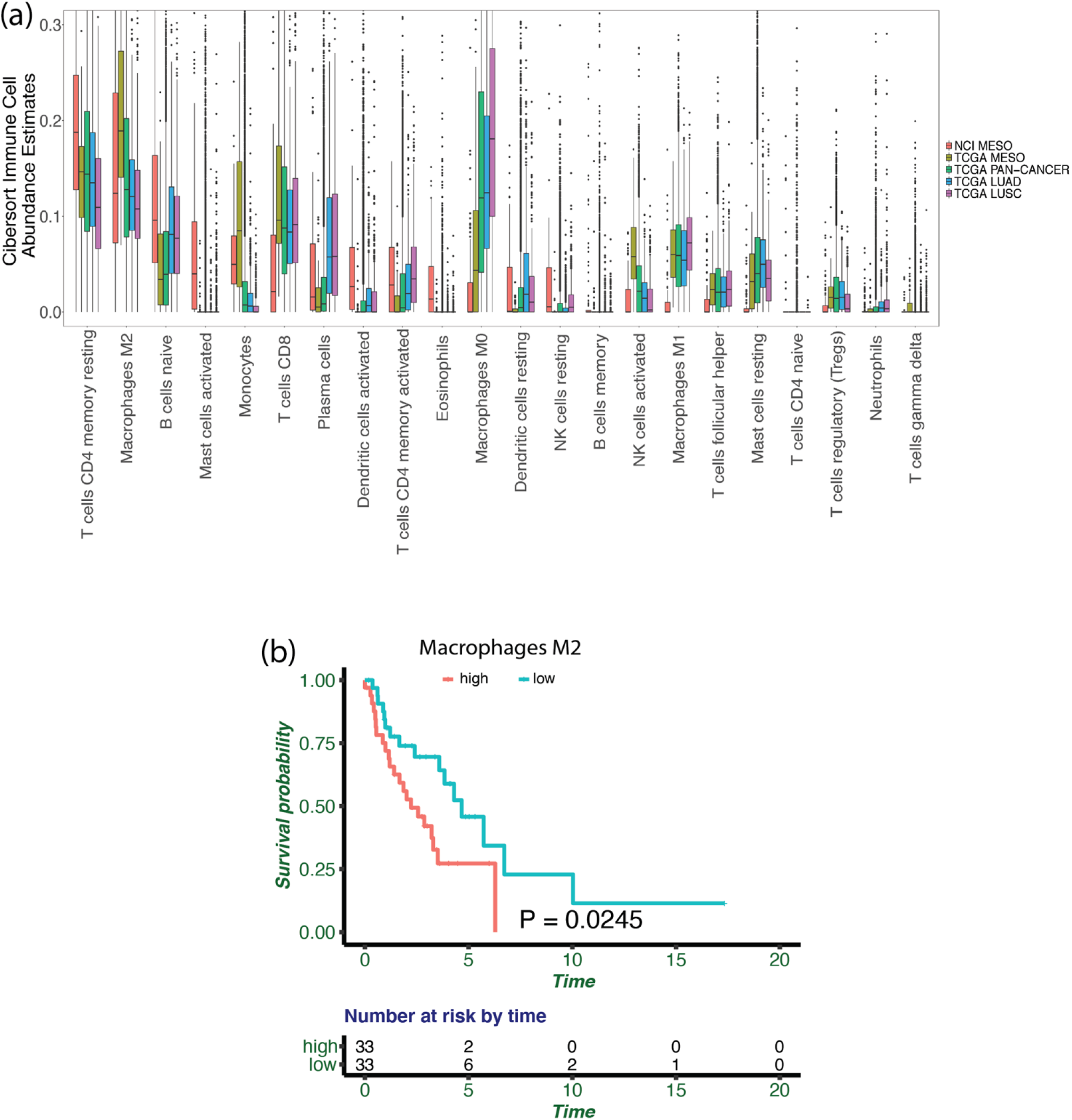
Immune cell abundance estimates. **(a)** Box plot of the relative fractions of the immune cell abundance across all mesothelioma samples in the NCI mesothelioma dataset (NCI MESO), TCGA pan-cancer patients (TCGA PAN-CANCER; except mesothelioma), TCGA lung adenocarcinoma (TCGA LUAD), TCGA lung squamous cell carcinoma (TCGA LUSC). The relative immune cell abundance for each immune cell type is shown as a fraction in the y-axis. **(b)** K-M survival plot of patients with top 33 and bottom 33 percentile of relative abundance of Macrophages M2 in the NCI mesothelioma dataset. Time in years. Log-rank test p-value is shown.

We tested the association of all the immune cell types that have more than 5% mean relative abundance across samples in the NCI mesothelioma dataset (Methods), with patient survival. We found that an increase in macrophages M2 was associated with worse mesothelioma patient survival in NCI mesothelioma dataset (Cox regression HR = 17.42, P=0.027, FDR = 0.16, after controlling for age and gender). We repeated this analysis using K-M survival plots and found similar results (**Fig. 4b**; log-rank P=0.0245). Since macrophages M2 are thought to be immunosuppressive (Noy & Pollard, 2014; Zhao et al., 2020), their association with decrease in survival was expected. We, however, did not see any association of macrophage M2 abundance estimates (using CIBERSORT) with patient survival in the TCGA mesothelioma and Bueno *et al*. datasets (FDR < 0.2). We also did not find any significant difference for any immune cell type abundance estimates between pleural and peritoneal mesothelioma patients (FDR < 0.2).

### A synthetic lethality/rescue transcriptome-based approach predicts responses to anti-PD1 therapy and combinations therapy with pemetrexed in patients with mesothelioma

Mesothelioma is difficult to treat cancer with very few therapeutic options available in the clinic. Therefore, a precision-medicine based approach for recommending therapies would be extremely useful. To this end, we employed the concept of synthetic lethality/sickness (SL) and synthetic rescue (SR) to predict patient response to therapies. Synthetic lethality is a genetic interaction between two genes, such that, when one of the genes is inactivated, the cells remain viable, however, when both genes are inactivated, the cells lose their viability (35,49). Synthetic rescue on the other hand is a genetic interaction between two genes such that, when any one of the genes is inactivated, the cells become less viable, but downregulation (or upregulation) of the partner gene rescues the cell (35,36,50). Lee *et al*. (2021) used a computational pipeline called SELECT (SynthEtic Lethality and rescue-mediated precision onCology via the Transcriptome), a precision-oncology approach, which aims to predict drug response for a given cancer patient using the whole pre-treatment transcriptome data. SELECT is based on inferring clinically relevant pan-cancer SL and SR pairs across cancer types by mining thousands of patient tumor genomic and transcriptomic data from the TCGA pan-cancer data and various *in vitro* studies [Methods, (35,36,50)]. For each chemotherapy or targeted-therapy drug, SELECT assigns a risk score based on the number of downregulated SL partners of the target genes inhibited by the drug. It works under the assumption that a drug is likely to kill a tumor more effectively when the SL partners of its targets are downregulated, as the inhibition of the targets by the drug will lead to the down regulation of the pertaining SL pairs and reduce cell viability and fitness. For immunotherapy drugs, SELECT assigns risk scores using SR interactions in an analogous manner (Methods). Using the SL/SR network, SELECT was previously shown to accurately predict drug response and stratify many patients in 28 of 35 independent targeted and immunotherapy datasets ranging across various cancer types and for many different drugs (36), using the same set of parameters. It is important to note that SELECT does not train any model parameters by looking at the test data, mitigating the risk of overfitting the data and loss of generalization predictive power on new, unseen datasets.

Among the 100 patients with expression data in the NCI mesothelioma data, 16 patients were treated with anti-PD1 immune checkpoint inhibitors (Pembrolizumab or Nivolumab, see Methods, **Tables S7a, S4b**). DrugBank was used to map drugs to targets that are inhibited (37). Similar to what was reported in other clinical trials (13), only a small fraction of these patients (18.8%; 3 out of 16 patients), responded to treatment (either complete or partial response). We used SELECT to predict patient response for anti-PD1 drugs in the NCI mesothelioma patient cohort, using the exact same parameters as used previously including the decision threshold value (36). Despite having a small number of responders, we were able to predict anti-PD1 drug response accurately in mesothelioma patients (ROC-AUC = 0.91; area under precision-recall or PR-AUC = 0.74; **Fig. 5a, Table S7a**).

**Figure 5:**
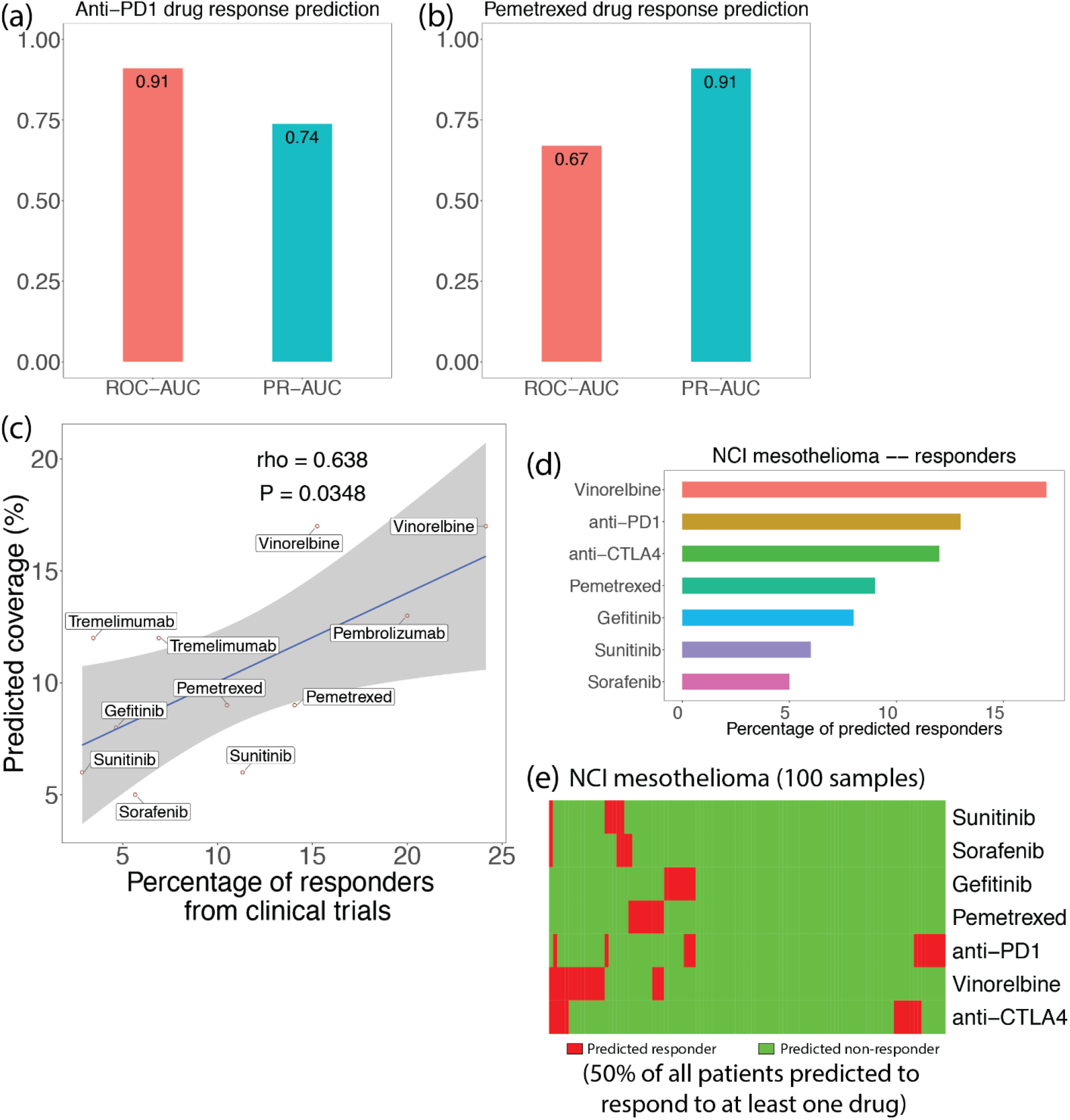
Overall SL transcriptomics based response prediction across several mesothelioma clinical trials. **(a)** We use SELECT to predict patient response for anti-PD1 drugs in the NCI mesothelioma patient cohort using patient gene expression data. Bar plots show ROC-AUC and area under precision recall (PR-AUC) value using the SELECT derived risk to predict responders (complete or partial response) from non-responders (stable disease or progressive disease) of anti-PD1 immunotherapy (16 samples). **(b)** Bar plots show ROC-AUC and area under precision recall (PR-AUC) value using the SELECT derived risk to predict responders (complete or partial response) from non-responders (progressive disease) of chemotherapy drug Pemetrexed (41 samples). **(c)** Scatter plot shows the percentage of responders (objective response rate) from mesothelioma clinical trials and comparing it with the percentage of predicted responders (coverage) using SELECT in the NCI mesothelioma dataset. Spearman’s ρ (rho) and p-values are shown. The shaded region is 95% confidence level interval for a linear model.**(d)** Bar plot showing the percentage of predicted responders for each drug. **(e)** Heatmap showing the responders in all patients and drugs is shown in NCI mesothelioma data. The x-axis corresponds to patient samples and y-axis is for each drug. Red color indicates the patient is predicted to respond to that drug, and green color indicates non-response. 50% of the total percent of patients are predicted to respond to at least 1 drug in the NCI mesothelioma dataset.

Pemetrexed is a commonly used chemotherapy drug, and it is often used in combination with other chemotherapy drugs like cisplatin or carboplatin. In the NCI mesothelioma cohort, pemetrexed was always given in combination with other drugs, and we considered the 41 patients who were given this drug (1 complete response, 23 partial response, 6 progressive disease, 11 stable disease; Methods, **Tables S7b, S4b**). We used the drug targets of pemetrexed as reported in Drugbank to predict its corresponding drug combination response, using SELECT. We could not consider the response to the other drugs (cisplatin/carboplatin) given in these combinations since they could not be mapped to targets that they inhibited according to DrugBank. Despite the limitations of this analysis, when we removed patients with stable disease from the analysis, we were able to predict patients with complete/partial response from those with progressive disease quite well (ROC-AUC = 0.67, PR-AUC = 0.91, **Fig. 5b, Table S7b**).

### SL-based transcriptomics based prediction of overall response to cancer drugs in numerous clinical trials in mesothelioma

We wanted to check if the concept of SL/SRs could be used to predict the percentage of responders to various mesothelioma cancer therapies. We focused on analyzing clinical trials using various targeted-therapies and immunotherapies, as these categories of drugs were previously shown to be well predicted by SELECT (36). We additionally considered a few chemotherapies commonly used in mesothelioma, like pemetrexed, vinorelbine, etc. We did not consider other commonly used chemotherapy drugs like cisplatin and carboplatin, as they could not be mapped to drug targets that they clearly inhibit (as per DrugBank). For each of the drugs, we collected information of the objective response rates (percentage of responders) in mesothelioma patients from many of the pertaining published clinical trials. We succeeded in obtaining this information for 7 drugs overall, including 2 chemotherapies, 3 targeted-therapies, and 2 immunotherapies: Pemetrexed, Vinorelbine, Gefitinib, Sorafenib, Sunitinib, Pembrolizumab, Tremelimumab (12-22) (**Table S7c**). The individual patients’ transcriptomic data was not available for these clinical trials so we could not aim to predict the individual response of each patient in these trials. However, we could still aim to study how well does the predicted SL-based coverage for each drug (based on the analysis of available mesothelioma patients’ transcriptome in our cohort) match the actual overall response observed in these trials.

To this end, SELECT was used to predict the percentage of responders (coverage) for each drug in the NCI mesothelioma cohort, using the same parameters and thresholds used (36) and repeating the exact same procedure that they have published (Methods). We found a significant correlation between the predicted and actual coverage of these drugs (Spearman’s ρ = 0.64, P = 0.0348, **Fig. 5c, Table S7c**). Notably, this also suggests that the NCI mesothelioma cohort may faithfully reflects the transcriptome of patients participating in these various clinical trials. For robustness, we performed the same analysis on TCGA mesothelioma patients and reassuringly, obtained similar results (**Fig. S7a**). **Fig. 5d**,**e** shows the fraction of patients predicted to respond to each of the 7 drugs considered, using SELECT (**Table S7d**,**e**). We found that 50% of the patients in the NCI mesothelioma data were predicted to respond to at least one of the 7 drugs based on the risk scores of the drugs computed for each patient (**Fig. 5e**). This percentage is higher for the TCGA data (64.71%). Since some patients were predicted to respond to more than one drug, we ranked the effectiveness of all drugs for each patient in the NCI mesothelioma cohort. Across the chemotherapies and targeted therapies considered, Vinorelbine is the highest ranked drug (see details in **Fig. S7b**) and this is in line with the outcomes observed in the clinical trials (21,22). Similar analysis was done for immune checkpoint inhibitors in mesothelioma (**Fig. S7c**).

## DISCUSSION

In this study we performed whole exome sequencing (WES) of blood and tumor tissues from 122 malignant mesothelioma patients who participated in our clinical trial, and systematically studied their somatic and germline mutations. We found that key somatic mutational signatures were associated with DNA repair pathways and *BAP1* was the most commonly mutated gene (∼13% with germline mutation). We also performed RNA-seq of 100 tumor tissues samples for transcriptomic analysis. This large-scale data resource along with the patient phenotypic characteristics and drug response information have been presented in a publicly available data resource. Our study cohort consists of patients with pleural (n=59) and peritoneal disease (n=61). To our knowledge, unlike previous studies which mainly focused on pleural mesothelioma patients, this is the first report that includes comparable number of patients of both MPM and MPeM subtypes, and hence gives us a better understanding of the similarities and differences that may exist in the molecular pathophysiology of the two anatomically distinct disease. Due to limited number of patients with tunica vaginalis mesothelioma, we are unable to compare the molecular characteristic for this disease.

As discussed in previous studies, we observe that the overall survival of patients with MPeM was higher than that of patients with MPM. Surprisingly, we did not find too many differences between pleural and peritoneal mesothelioma samples at the level of germline or somatic mutations, or at the level of copy-number information.

We further stratified the patients based on their overall survival and analyzed the differentially expressed genes in the high and low risk subgroups. We identified a “mesothelioma prognostic signature”, a set of 48 genes, that were overexpressed in high-risk cohort with poor survival and showed that the median expression of these genes is highly predictive of mesothelioma patient survival in two independent large-scale patient cohorts, and thus may have high translational value in the clinic. We found that these genes were enriched for GO terms related to cell cycle processes, and DNA repair, as well as others. Among these 48 genes, we found that the *CCNB1* gene alone was highly predictive of mesothelioma patient survival.

PPI of these 48 mesothelioma prognostic signature genes identified 3 cliques and two protein complexes including *CCNB1* and *CCNF* genes. This complex was not identified in other studies (10). Two genes in the signature, *CCNB1* and *NUSAP1* gene were found to be differentially upregulated genes in mesothelioma cancer compared to normal samples in previous literature. A transcription factor enrichment analysis also showed that that *CCNB1* gene is potentially regulated by other TFs. Cyclin B1, a protein encoded by the *CCNB1* gene, is a key mitotic cyclin in the G2-M phase transition of the cell cycle and is overexpressed in various malignant tumors, including lung, breast, colorectal, pancreatic, and others. Previous work has shown that Cyclin B1 plays an important role in tumor development and tumorigenesis (51,52). A meta-analysis about the prognostic role of cyclin B1 in solid tumor was conducted on 17 published studies, which made a conclusion that overexpression of cyclin B1 is a significant prognostic parameter and is associated with poor survival in many solid tumors (53). However, to the best of our knowledge there is no prior work showing that the expression of *CCNB1* gene by itself is associated with poor survival of mesothelioma patients, and this is the first study to do so. *CCNF* gene which encodes Cyclin F, is also a member of the cyclin family, is known to control genome instability and may play a role in cancer development (54). *NUSAP1* gene is a nucleolar-spindle-associated protein that it is known to play a role in spindle microtubule organization (55) and is known to play a role in some other cancers (56-58) but has not been significantly discussed in mesothelioma. Looking at CRISPR-Cas9 based essentiality screens in pleural mesothelioma cell lines, we found that the mesothelioma prognostic signature genes tend to be more essential; and we found about 10 genes which were highly essential (including the *MTBP* and *CCNB1* genes). *MTBP* gene is known to play a role in cell migration and invasion in some tumors (59-61), and to our knowledge has not been discussed in mesothelioma. Thus, using various analysis, we were able to identify a subset of the 48 genes in the mesothelioma prognostic signature which are likely to play an important role in mesothelioma. We however note that our studies are associative, and further *in vitro* and *in vivo* work is required to show any causal relationships for any of the mesothelioma prognostic signature genes to mesothelioma cancer progression.

Anti-tumor immune response for tumor clearance is utilized by tumor antigen direct immunotherapy and use of check point inhibitors. Hence it is translationally relevant to understand the tumor microenvironment. We used CIBERSORT to determine immune cell lineages infiltrating the tumor tissue based on immune cell specific marker expression and compared the immune abundance estimates in mesothelioma with other cancer types. Ideally, we would like to compare mesothelioma immune abundance with the corresponding mesothelial tissues but given the absence of large-scale normal mesothelial tissue expression data; we simply compared the relative immune estimates of the NCI mesothelioma dataset with the relative immune estimates in the TCGA cancer dataset. We found that some immune cell types like T cells CD4 memory resting cells, B cells naïve, etc. are higher relative abundance in NCI mesothelioma data than some of the other cancer types in the TCGA. We corelated the gene expression data with patient survival and found that increased tumor macrophage M2 was associated with poor survival. This is not surprising, since M2 macrophages are immunosuppressive in function.

A precision-oncology based approach called SELECT was shown to effectively predict drug response for various cancer treatments across many different cancer patients and cancer types, using patient transcriptomic data and the concepts of synthetic lethality and rescues (36). Using exactly the same parameters and thresholds used by (36), we applied SELECT on the NCI mesothelioma dataset, and were able to effectively predict anti-PD1 immune checkpoint inhibitor response. We were also able to predict drug combination response for an important chemotherapy drug like pemetrexed. Using SELECT to predict the percentage of responders (coverage) for various drugs in the NCI mesothelioma cohort (and also in TCGA mesothelioma) for which we know the response rate based on mesothelioma clinical trials, we found a significant correlation between the predicted and actual coverage for these drugs. This suggest that the NCI mesothelioma cohort may have similar transcriptomic heterogeneity seen in various clinical trials. As we did not expect synthetic lethality or rescue would explain the variation of drug coverage for all treatments, it was indeed surprising that we can explain the variation in patient response to different commonly used cancer therapies at a such high degree. As far as we know, this is the first factor to predict drug response and coverage for various treatments within mesothelioma. Using SELECT, we were able to predict effective treatments for about 50% of the patients in the NCI mesothelioma dataset and rank drugs based on effectiveness for each patient. Obviously, the veracity of these predictions needs to be studied carefully in controlled prospective clinical trials in the future.

The SELECT based prediction analysis performed has a few limitations that are important to note. First, the number of patient samples for drug response analysis for the anti-PD1 checkpoint inhibitors and pemetrexed is small, and the responding vs non-responding classes are imbalanced. Secondly, we attempted to predict drug combinations involving pemetrexed using only the drug targets of pemetrexed, since pemetrexed was only given in combination with other drugs. Moreover, some of the patients were previously treated with other various treatments. Thirdly, for some of the drugs for which we aimed to predict the overall response in clinical trials, we did not have drug response and transcriptomic information in mesothelioma cohorts. Therefore, further drug response validations in large scale mesothelioma patient cohorts are required to evaluate the value of such precision oncology approaches. Given that mesothelioma is a very difficult cancer to treat, we think that our precision-oncology framework provides a proof of concept to motivate further experimental studies and clinical trials in mesothelioma.

In conclusion, by analyzing genome and tumor transcriptome of patients with pleural and peritoneal mesothelioma, we have been able to identify deregulated pathways and prognostic markers that can be therapeutically targeted, and gene expression profiles predictive of patient survival and tumor response to available therapies.

## DATA AVAILABILITY

The NCI mesothelioma data resource is made publicly available in https://clinomics.ccr.cancer.gov/clinomics/public/ website, dbGaP (accession number: phs002207) and Table 1, Tables S1-S4.

## ACKNOWLEDGEMENTS

This research was supported in part by the Intramural Research Program of the National Institutes of Health (NIH), National Cancer Institute and the Center for Cancer Research. This work utilized the computational resources of the NIH HPC Biowulf cluster (http://hpc.nih.gov). The results shown here are in part based upon data generated by the TCGA Research Network: https://www.cancer.gov/tcga. We would like to thank Dr. Kuoyuan Cheng for this help on this project.

