## Supplementary notes and supplementary figures for "Transcriptomics based prediction of survival and response to therapy in malignant mesothelioma"

**SUPPLEMENTARY APPENDIX**

**Supplementary Results**

**Supplementary Figures (Figures S1-S7)**

**Supplementary Tables (Tables S1-S7) (Excel Sheets)**

**Table S1:** Occurrence of germline pathogenic and likely pathogenic variants in NCI mesothelioma cohort

**Table S2:** Copy number information for 122 patients in the NCI mesothelioma dataset

**Table S3:**

**a)** Gene expression data in the NCI mesothelioma dataset for 100 patient samples that we used in our differential expression analysis. Count data is shown.
**b)** Gene expression data in the NCI mesothelioma dataset for 100 patient samples that we used in our analysis. The data was in TMM+log(CPM) format, normalized using edgeR package. Rows which could not be mapped from Ensembl to common gene names were removed. This data was used for our survival analysis.

**Table S4:**

**a)** Phenotypic information for various patients in the NCI mesothelioma dataset.
**b)** Drug response information for many of the NCI mesothelioma patients.

**Table S5:**

**a)** Genes in mesothelioma prognostic signature. These are 48 genes whose increased expression is associated with decreased survival (FDR < 0.1). Using Cox regression after controlling for age, sex, and site of disease.
**b)** Genes whose increased expression is associated with increased survival (FDR < 0.1). Using Cox regression after controlling for age, sex, and site of disease.
**c)** GO enrichment analysis on mesothelioma prognostic signature genes. Using GOrilla tool on 'Process' ontology.
**d)** Risk scores for NCI mesothelioma dataset using median gene expression of mesothelioma prognostic signature.
**e)** Genes whose increased expression is associated with decreased survival (FDR < 0.1). Using Cox regression after controlling for age, sex, and site of disease.
**f)** GO enrichment analysis on overlapping genes. Using GOrilla tool on 'Process' ontology.
**g)** PPI network (clique 1) protein complex enrichment analysis. https://mips.helmholtz-muenchen.de/corum/#download. 03.09.2018 Corum 3.0 current release.
**h)** Transcription factor regulating genes whose increased expression is associated with decreased survival (FDR < 0.1, mesothelioma gene signature). We used TRRUST version 2 website for this analysis.
**i)** Essentiality values for 7 pleural mesothelioma cell lines (CRISPR-Cas9 gene knockout data from Doench et al., 2016; Meyers et al., 2017) and also median essentiality. Data is shown for 44 out of the 48 genes in the mesothelioma prognostic signature.

**Table S6:** Immune abundance estimates using CIBERSORT in the NCI mesothelioma dataset. CIBERSORT was run for 500 permutations using the default gene signature LM22 CIBERSORT, from the bulk tumor gene expression data. The results were run in the ‘Relative’ mode generating ‘Relative’ fractions of 22 immune cell types for each of the 100 patients in the NCI mesothelioma dataset. The relative values indicate the relative abundance of each immune cell type.

**Table S7:**

**a)** Patients from the NCI mesothelioma data that were considered for anti-PD1 drug response analysis. Patient response and predicted risk scores using SELECT are provided.
**b)** Patients from the NCI mesothelioma data that were considered for Pemetrexed drug response analysis. Patient response and predicted risk scores using SELECT are provided.
**c)** Clinical trials considered for mesothelioma cancer patients, their response rates, and their predicted coverage.

**d)** SELECT risk score predictions for chemotherapy and targeted-therapy in NCI mesothelioma dataset patients. Higher the score, more likely the patient will respond to a drug. Risk scores > 0.44 are predicted to be responders.

**e)** SELECT risk score predictions for immunotherapy in NCI mesothelioma dataset patients. Higher the score, more likely the patient will respond to a drug. Risk scores >= 0.9 are predicted to be responders.

### SUPPLEMENTARY RESULTS

#### Mesothelioma prognostic signature

We identified 48 genes in the NCI mesothelioma data, whose increased expression is associated with worse survival using Cox regression analysis, after controlling for age and sex, with FDR < 0.1. We call this set of 48 genes as "mesothelioma prognostic signature" gene-set. These 48 signature genes are: *TRIP13, MCM2, MCAM, UBE2T, KIF4A, MDK, FOXM1, COL7A1, CIT, CKS2, CCNB1, COL4A2, CCNJL, TROAP, NUSAP1, SMAD6, CDH24, CDCA5, INCENP, EME1, HIST1H4H, RECQL4, RACGAP1, CCNF, PAQR4, EDIL3, RCOR2, GPRIN1, ALCAM, MTBP, CSPG4, UBE2C, FZD2, OLFML2A, KIF18B, COL4A1, HIST1H2AI, MAFG, IPO9, PRC1, MZT1, AJM1, KIFC1, HOXA10, HIST1H4K, HIST1H3A, HIST1H4E, HIST1H2AE.*

#### Enrichment analysis of survival genes across datasets

We checked the association of gene expression with survival for TCGA mesothelioma and Bueno *et al.* datasets separately, using Cox regression after controlling for age and sex. For the TCGA dataset, there are 2144 genes whose increased expression is significantly associated with decreased survival and 2181 genes whose increased expression is significantly associated with increased survival (FDR < 0.1). For the Bueno *et al.* dataset, there are 1210 genes whose increased expression is significantly associated with decreased survival and 1062 genes whose increased expression is significantly associated with increased survival (FDR < 0.1).

We take the intersection of genes whose increased expression is associated with decreased survival (FDR < 0.1) in all three datasets – NCI mesothelioma, TCGA mesothelioma and Bueno *et al.* mesothelioma datasets. We find that there are 31 genes which show this pattern in all three datasets (**Table S5e**). These 31 genes may be strong candidates for drug targets. For the genes which are associated with survival (FDR < 0.1) in each of the three datasets, we tested if the overlapping genes have an enrichment (using hypergeometric test). Between the NCI mesothelioma dataset and TCGA mesothelioma dataset, we find that there is a significant overlap (intersection) of survival associated genes (P = 9.41e-15). Between the NCI mesothelioma dataset and Bueno *et al.* mesothelioma dataset, we find that there is a significant overlap (intersection) of survival associated genes (P = 1.85e-14).

#### Alternative method of computing risk

For the survival analysis, we also computed the risk scores using an alternative method. The new *fractional risk score* for a patient was computed by counting the fraction of the 48 genes in the mesothelioma prognostic signature which are highly expressed (top 33 percentile of all genes) in a patient. Survival analysis for the new risk scores is done on TCGA and Bueno *et al.* mesothelioma datasets after controlling for age and sex (see **Fig. S5**). We see that the results are quite similar to the previous method of computing combined risk scores by taking median expression of the signature genes (i.e., analysis in **Fig. 2b**). This shows that our mesothelioma gene signature is quite robust.

**SUPPLEMENTARY FIGURES**

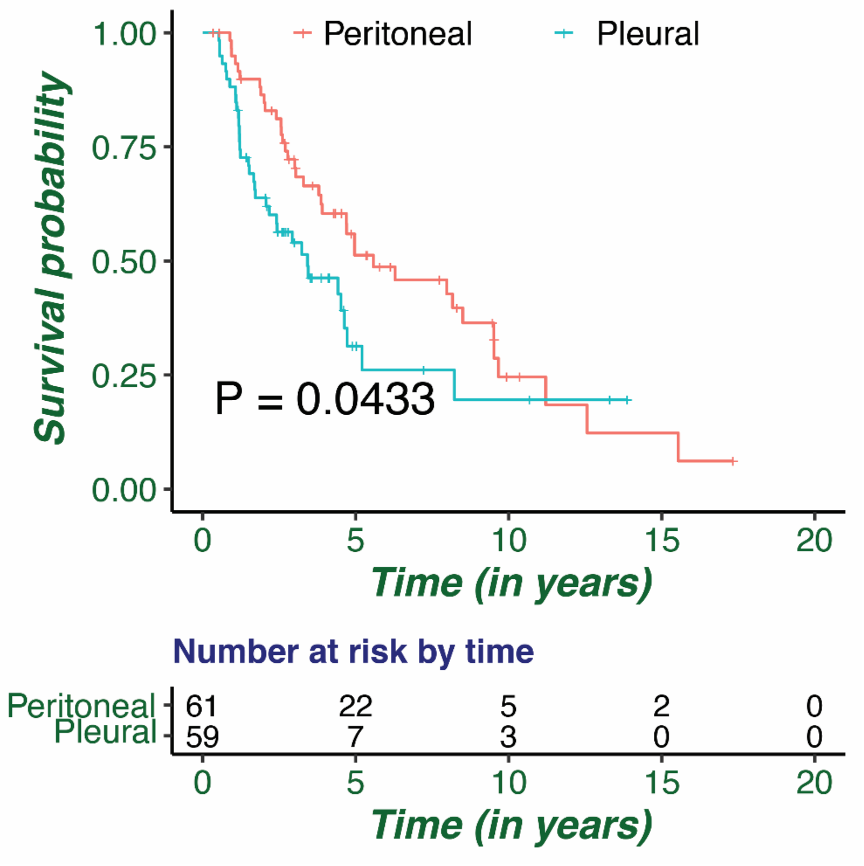


***Figure S1:*** *K-M plot showing overall survival differences between Pleural and Peritoneal mesothelioma. Log-rank test p-value between the two groups is shown.*

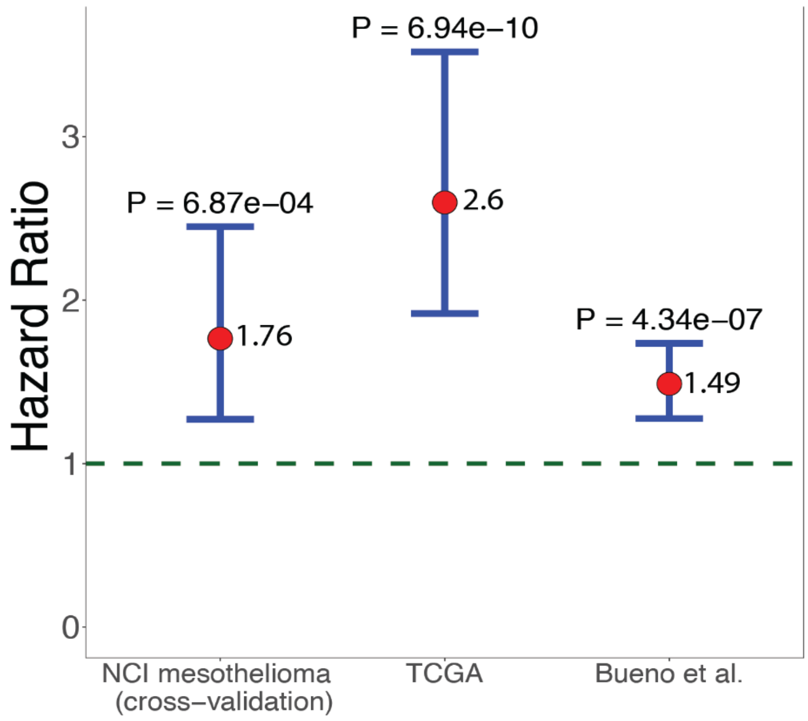


***Figure S2:*** *Cox regression results of predicted risk score and mesothelioma patient survival in TCGA and Bueno et al. datasets after controlling for age and sex. The Cox regression results for NCI mesothelioma dataset is shown as cross-validation, after controlling for age, sex, and site-of-disease. Plots were computed using the median values of 48-gene high risk/low survival mesothelioma prognostic signature genes. Hazard ratios (HRs) greater than 1 indicate increased risk and is associated with worse survival.*

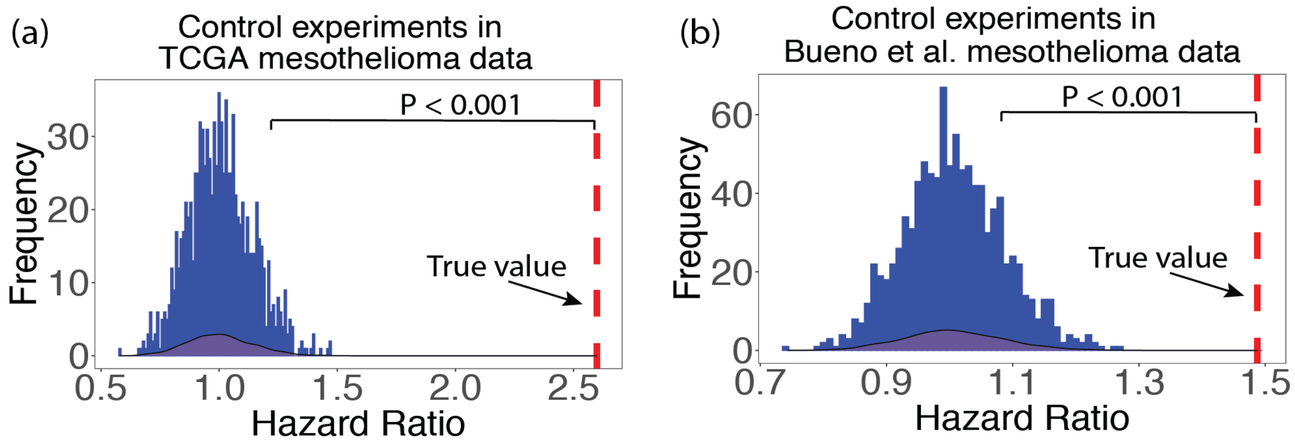


***Figure S3:*** *Control experiments predicting TCGA mesothelioma and Bueno et al. mesothelioma datasets using the random set of genes of comparable size to those of mesothelioma gene signature genes. The random sampling was done for 1000 iterations. The randomization test P-value is shown with the Cox regression Hazard ratios of the random controls with that obtained using the mesothelioma gene signature.*

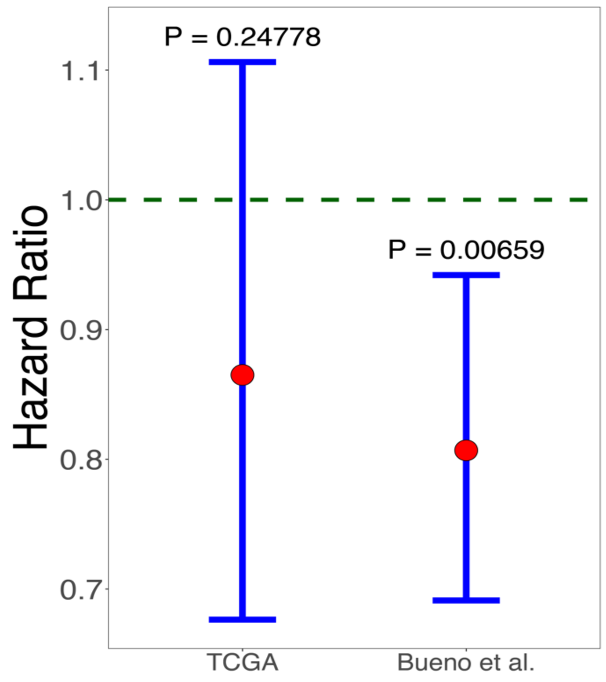


***Figure S4:*** *Cox regression survival analysis results of predicted risk score in TCGA and Bueno et al. mesothelioma datasets after controlling for age and sex. Median expression of genes whose increases expression is associated with increased mesothelioma patient survival was used to calculate risk score.*

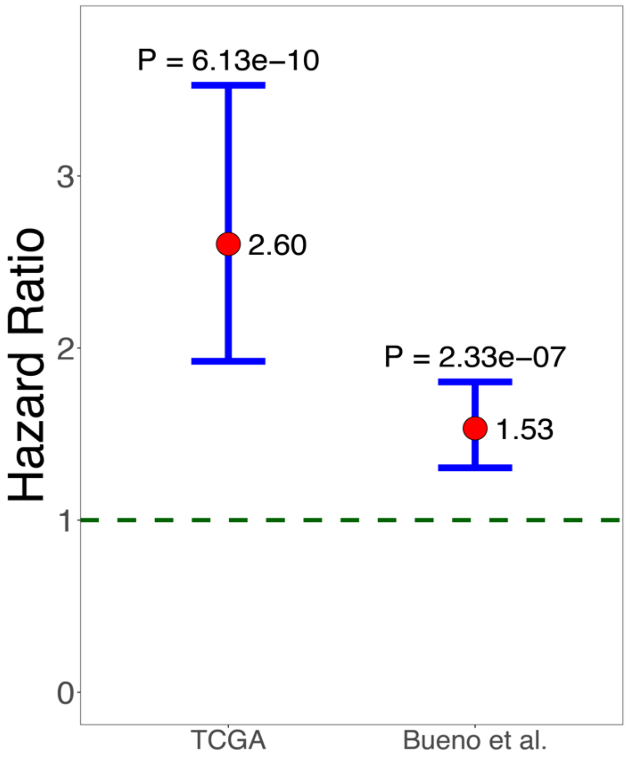


***Figure S5:*** *Cox regression survival analysis results of predicted fractional risk score using an alternative method on the mesothelioma gene signature (48 genes). Risk score for a patient was computed by counting the fraction of the 48 genes in the mesothelioma prognostic signature which are highly expressed (top 33 percentile of all genes) in a patient. Survival analysis for the new risk scores was performed on TCGA and Bueno et al. mesothelioma datasets after controlling for age and sex.*

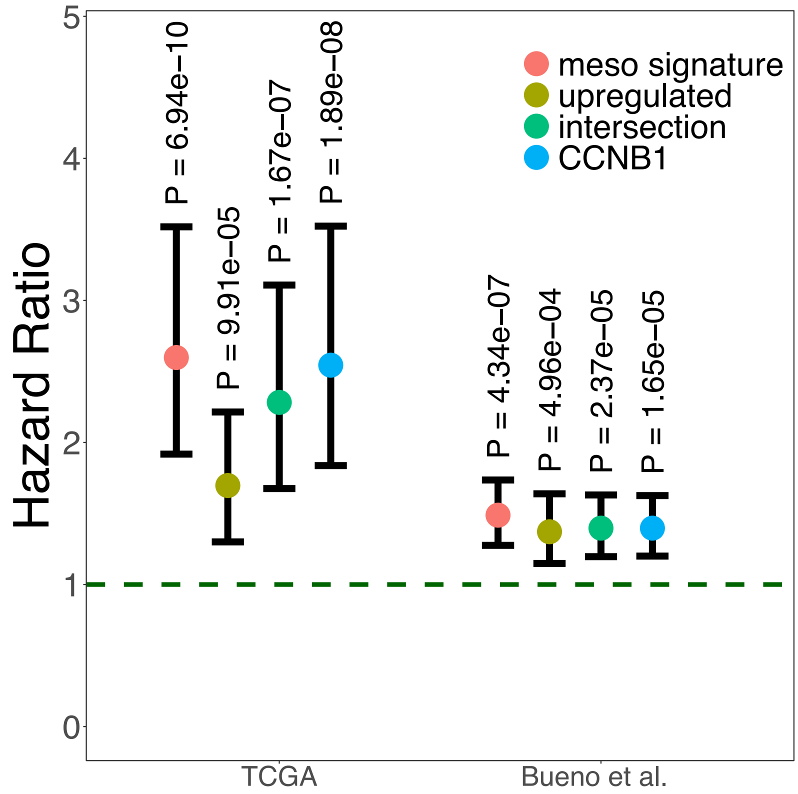


***Figure S6:*** *Cox regression of predicted risk score with mesothelioma patient survival in TCGA and Bueno et al. datasets after controlling for age and sex. Risk scores were predicted using various methods: median expression of 48-gene (high-risk/low-survival) mesothelioma prognostic signature (signature); median expression of 51 differentially upregulated genes in mesothelioma cancer compared to normal samples in independent studies (upregulated); median expression of 2 genes (CCNB1, NUSAP1) which overlap between the 48-gene mesothelioma prognostic signature and 51 differentially upregulated genes; using gene expression of only CCNB1 gene. Hazard ratios greater than 1 indicates increased risk associated with worse survival.*

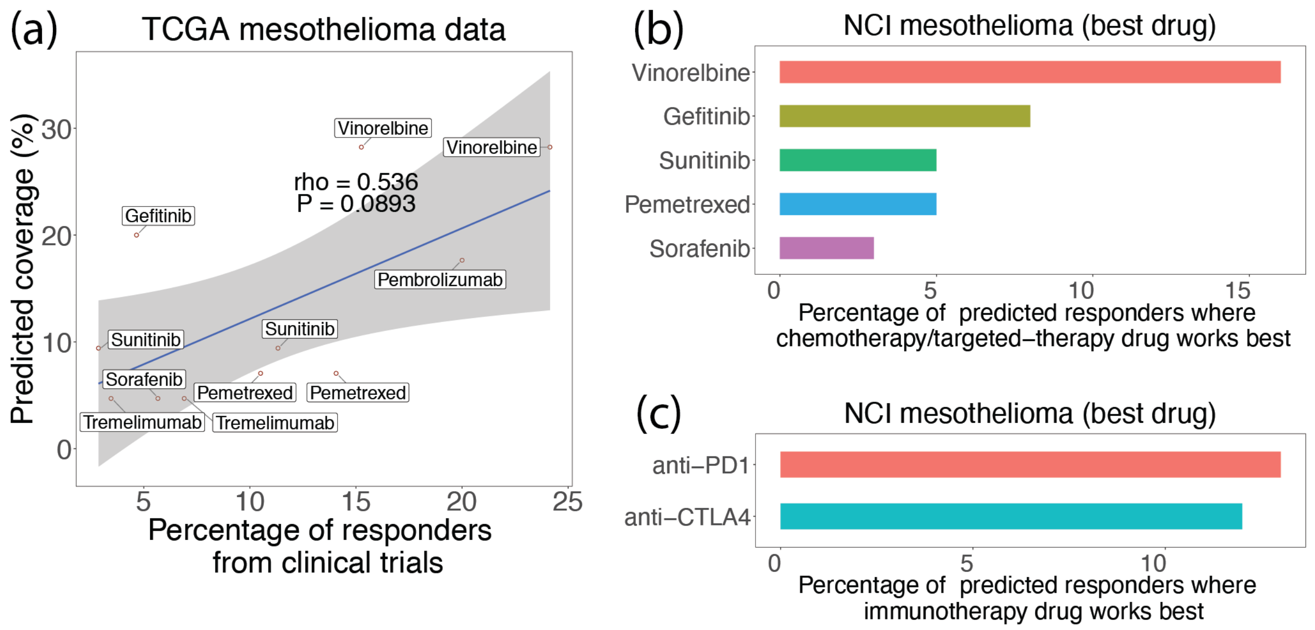


***Figure S7:*** *(a) Scatter plot shows the percentage of responders (objective response rate) from mesothelioma clinical trials and comparing it with the percentage of predicted responders (coverage) using SELECT in the TCGA mesothelioma dataset. Spearman's ρ (rho) and p-values are shown. (b) For each patient in the NCI mesothelioma dataset, we rank all chemotherapy or targeted therapies we considered based on their effectiveness. For each drug, we plot the percentage of NCI mesothelioma patients where this drug worked best among all other drug, provided the predicted risk score > 0.44 (shown as a bar plot). (c) For each patient in the NCI mesothelioma dataset, we rank all immunotherapies we considered based on their effectiveness. For each drug, we plot the percentage of NCI mesothelioma patients where this drug worked best among all other drug, provided the predicted risk score ≥ 0.9 (shown as a bar plot).*
